# Temporal Gene Coexpression Network Analysis Using A Low-rank plus Sparse Framework

**DOI:** 10.1101/359612

**Authors:** Jinyu Li, Yutong Lai, Chi Zhang, Qi Zhang

**Author notes:** These two authors contributed equally to this work (Jinyu Li, Yutong Lai).

## Abstract

Various gene network models with distinct physical nature have been widely used in biological studies. For temporal transcriptomic studies, the current dynamic models either ignore the temporal variation in the network structure or fail to scale up to a large number of genes due to severe computational bottlenecks and sample size limitation. On the other hand, correlation-based gene networks are more computationally more affordable, but have not been properly extended to gene expression time-course data.

We propose Temporal Gene Coexpression Network (TGCN) for the transcriptomic time-course data. The mathematical nature of TGCN is the joint modeling of multiple covariance matrices across time points using a “low-rank plus sparse” framework, in which the network similarity across time points is explicitly modeled in the low-rank component. Using both simulations and a real data application, we showed that TGCN improved the covariance estimation loss and identified more robust and interpretable gene modules.

## 1 Introduction

High throughput manner and low cost of sequencing technologies enabled the biologists generating an enormous wealth of data for discovering and quantifying the relationship among large amounts and various types of molecular elements, such as gene expressions, proteins, metabolites and epigenetic marks. These elements and their relationship or interactions could be modeled as nodes and edges in a network model. Specifically, the gene co-expression network (GCN) models have been used for the exploration, interpretation and visualization of the relationship among genes in a wide range of biological applications, including disease-gene association (Yang *et al*., 2014), identification of genes responding to environment changes, tissue specific gene identification (Tan *et al*., 2017), and functional gene annotation (Kadarmideen and Watson-Haigh, 2012). GCN outputs can also be combined with other biological data in various downstream analysis, such as identifying functional eQTLs (Villa-Vialaneix *et al*., 2013) and studying gene-phenotype association (Ficklin *et al*., 2010). Partially for this reason, many GCN databases were developed as functional annotation resources (e.g., GeneFriends van Dam *et al*. 2014, COXPRESdb Obayashi *et al*. 2007 and PlaNex Yim *et al*. 2013).

Many construction tools and analysis tools were developed for GCN, and the physical nature of the resultant networks are different. WGCNA (Zhang and Horvath, 2005) is based on the marginal correlation of the gene pairs. GeneTS (Schafer and Strimmer, 2004) and BicMix (Gao *et al*., 2016) built Gaussian Graphical Models (GGM) of genes, which were based on partial correlations. Under the multivariate Gaussian assumption, GGM captures the conditional dependence among genes. Mutual information has also been used in defining gene networks (Daub *et al*., 2004). Comparing with Pearson’s correlation, it could also capture the nonlinear association (Song *et al*., 2012). Different from the undirected networks produced by the above methods, Bayesian Network (BN) based methods infer directed networks from gene expression data (Friedman *et al*., 2000; Ni *et al*., 2015).

Temporal transcriptomic data are extremely useful in developmental biology(Hawrylycz *et al*., 2012) and stress biology(Miao *et al*., 2017), and GCN models have been utilized for such analysis. For instance, BNs could also be extended as Dynamic Bayesian Network (DBN) for time course data, which models the directed gene-gene relationship across time points (Lebre *et al*., 2010). Besides DBN, another line of dynamic network models are based on differential equations (Zhang *et al*., 2018). The major bottlenecks in popularizing these dynamic network models lie in the nature of the inference goal and the biological nature of the data. First, these computational algorithms search in high-dimensional parameter space and require large number of replicates, even for a small network. For example, Zhu and Wang (2015) reported that it took nine minutes for their proposed method HMDBN to learn a simulated dynamic network among 10 nodes using 1,019 observations, while its competitors all took 11 to 58 hours. Even though HMDBN has made tremendous progress along this direction, it may be still unrealistic to fit large DBN without further computational improvement, as the parameter space grow exponentially with the number of nodes. Secondly, for the models applicable to smaller samples (Oates and Mukherjee, 2012; Zou and Conzen, 2004), the structure of DBN are often assumed to be time-invariant, which could be unrealistic for the biological systems studies in developmental biology (Hawrylycz *et al*., 2012) and agronomy (Xu *et al*., 2012).

Due to the limitations and challenges in dynamic network modeling, biologists usually adopt one static GCNs to aggregate all data together or construct several static GCNs to analyze transcriptomic data at different time points (Ballouz *et al*., 2015; Miao *et al.*, 2017). Such strategies either mixes various sources of heterogeneity or could not describe the dynamics of the biological systems, but both make the interpretation of the resulting network difficult and inaccuracy. Alternatively, one may build one GCN using the data at each time point, and compared them using differential network analysis (Azevedo *et al.*, 2017). This approach also has many drawbacks. First, the replicates at each time point may not be large enough to build a reliable and robust GCN. Second, ignoring the similarity of the gene expression at different time points within the same time-course often results in false positives in differential network analysis across time points. In this manuscript, we presented a network model - the Temporal Gene Coexpression Network (TGCN), a new “low-rank plus sparse” framework (Fan *et al*., 2013; Luo, 2011) for building time-point specific GCNs from time-course gene expression data. In both simulation studies and the real data application, we showed that the resulting GCN’s were more robust and accurate than the separate modeling at each time point, and thus led to more accurate downstream analysis.

## 2 Methods

### 2.1 Model setup and notations

In a typical transcriptomic time course study, suppose we collect *n*_*t*_ samples, and observe the *p* × *n*_*t*_ data matrix *X*_*t*_ = (*x*_*t1*_,…, *x*_*t,nt*_) for *t* ϵ {*t*_1_,…, *t*_*T*_}, where p is the number of genes, *n*_t_ is the number of samples and *>T*> is the total number of time points. Reconstructing the GCN at each time point can be achieved by estimating *Σ*_*t*_, the covariance matrix of the columns of *X*_*t*_. Since this is our focus, we assume the rows of *X*_*t*_ have zero mean.

One naive estimate of *Σ*_*t*_ is simply the raw sample variance-covariance matrix 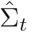. This approach may lead to extremely noisy estimates of GCN, because the sample size at each time point is usually very small. It also ignores the natural partial ordering of the samples (by time), and the similarity in the covariance structure among them. Such similarity comes from various sources. For example, the genes in the same pathway tend to be coexpressed, and the membership of the genes to the pathways are fixed. The temporal change in GCN, however, could be caused by the fact that these pathways share genes, and their activity intensity also vary through time.

The above biological observation motivates us modeling GCN with time-invariant latent factors, and their time-varying loadings, an approach connected with the low-rank approximation of high-dimensional covariance matrix (Fan *et al*., 2008). In the literature of matrix recovery, it has been noted that the low-rank approximation may be too restrictive and not robust, and a natural extension is the “low-rank plus sparse” framework (Fan *et al*., 2013; Luo, 2011). Towards this end, we also included a time specific sparse component to reserve the significant links that cannot be captured by the factor model.

To summarize, we propose the following “low-rank plus sparse” estimator for Σ_*t*_

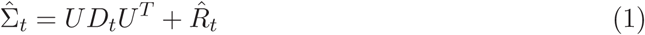

Here *U* is a *p* × *K* matrix whose columns are the time-invariant latent factors learned from the transcriptomic data itself, the *K* × *K* diagonal matrix D_t_ are their time-varying loadings, and 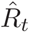 denotes the estimated sparse component of the time-specific links at time point *t*.

Throughout this paper, we will use *A(i,j)* to denote the element of *A* in its ith row and *j*th column, and (*b*)_+_ to denote the positive part of *b*, i.e. it is 0 if *b* is negative.

### 2.2 TGCN: Temporal Gene Coexpression Network analysis

We propose Temporal Gene Coexpression Network (TGCN, Algorithm 1) framework based on the low-rank plus sparse model Equation (1).

**Algorithm 1.**
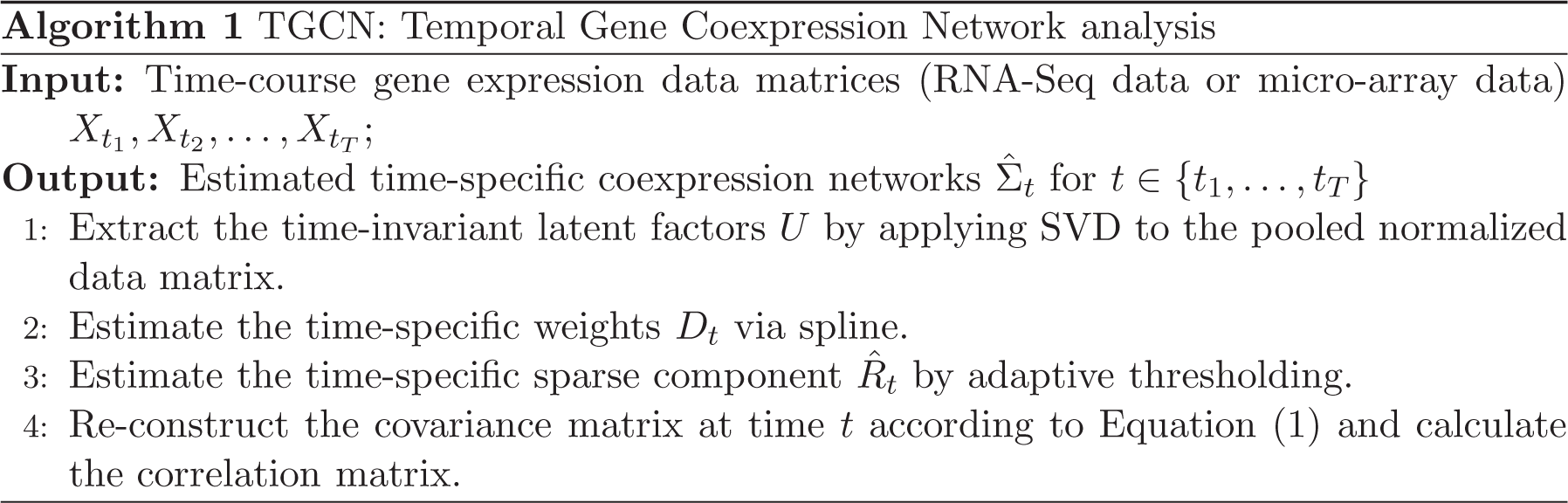
TGCN: Temporal Gene Coexpression Network analysis

When performing SVD on pooled data for extracting U in the first step, normalization is critical. If not done properly, the leading principle directions could be heavily influenced by the time points with larger sample sizes or larger variability. Data normalization typically involves centering and scaling. Our goal in this step is to find the principle directions that are representative at all time points. Thus we scaled Xt with the leading singular value of *X*_*t*_ and then multiply the square root of number of genes 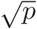. This approach assigned equal weights to the principle directions of the data at each time point. Write the pooled normalized data as

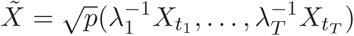

where λ_*l*_ is the leading singular value of *X*_*tl*_. Then we applied SVD to 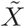, and used its top *K* left singular vectors as the columns of the time-invariant latent factors matrix U. Choosing the rank for low-rank matrix approximation is a difficult task (Valle *et al*., 1999), because fewer PCs will include incomplete information of the process while more PCs will cause the model over-parameterized and include noise. This problem could be posed as a model selection problem, and many information criteria have been applied, including Akaike’s entropy-based Information Criterion (AIC, Akaike 1973, 1998) and Minimum Description Length (MDL, Rissanen 1978, 1983; Valle *et al.* 1999. See Supplementary Notes for details). There is no dominant procedure for this problem, and both AIC and MDL are consistent under certain regularity conditions. However, Valle *et al.* (1999) suggested that AIC tended to overestimate the dimension of the low rank approximation in certain scenarios. This is consistent with our data analysis, where we found that both of AIC and MDL selected the correct number of components in simulations (Supplementary Fig. 1), while MDL selected a more compact model in the real data analysis (Supplementary Fig. 2). Thus we recommended using MDL at this step.

In the second step, one natural raw estimate of the weights of the latent factors U at time point t is the diagonal elements of 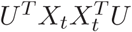. Let this length *K* vector be 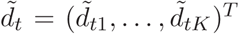. For *k* = 1,…, *K*, we propose to further smooth this raw estimates 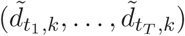 at its log scale using spline, and report resultant sequence (*d*_*tl*_, _*k*_,…, *d*_*tT*_, _*k*_) as the time-varying weights of this factor. Finally, we define

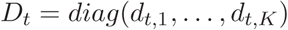

In the third step, we estimated the time specific sparse component 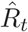 using the adaptive thresholding rule (Cai and Liu, 2011) as used in Fan *et al.* (2013). In detail, let

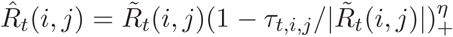

be the original residual matrix after removing the low-rank component from the sample covariance at t.The elements of the sparse component matrix are then

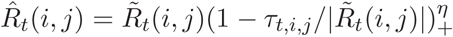

for 1 ≤ *i,j* ≤ *p* and *i* ≠ *j*, where the adaptive threshold

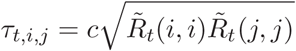

for some non-negative constant c € [0,1] and η > 0. In our analysis, we set c = 0.5, and η = 4 as suggested in Cai and Liu (2011) and Fan *et al.* (2013).

Finally, the time-specific gene-gene correlation matrices are obtained by normalizing the covariance matrices obtained from Equation (1).

### 2.3 Downstream Analysis of TGCN

The output of TGCN are time-specific gene-gene correlation matrices which could serve as the input of gene co-expression network analysis procedures such as *weighted correlation network analysis (WGCNA)* (Langfelder and Horvath, 2008). WGCNA is a method for finding clusters/modules of highly correlated genes, and then describing the correlation patterns among genes across different samples. In this study, we fed our outputs to WGCNA as adjacency matrices for constructing scale-free gene networks at each time point using power transformation and for module discovery by hierarchical clustering with adjacency-based dissimilarity.

In the real data analysis, we further investigated the biological interpretation of the discovered modules using R/Bioconductor package `clusterProfiler` (Yu *et al*., 2012). `clusterProfiler` performs hypergeometric test for enrichment analyses of given gene lists. We only performed the enrichment analysis of KEGG pathways instead of the other gene oncology terms, as we only focused on the genes that are associated with KEGG pathways.

## 3 Results

In this section, we investigate the properties of TGCN, and compared it with the gene coexpression networks constructed separately at each time point. We refer to this method as the “Naive” estimate as it ignores the association between time points.

## 3.1 Simulation-based evaluation

### 3.1.1 Simulation Model

We simulated gene expression data whose covariance matrices *Σ*_*t*_ were generated based on Equation (1). Let p be the number of genes, *C* the number of latent factors. Roughly speaking, the larger *C* is, the more complex the covariance matrices are. We simulated the time-invariant latent factor matrix *U* in a way that it encoded *C* potentially overlapping gene groups, and each factor was associated with a smooth weight curve. The time point specific sparse components *R*_*t*_’s were generated such that they also contained the same grouping structure. The transcriptomic time-course data were then generated from *N*(0, *Σ*_t_) (See Supplementary Notes for details of the simulation model). Let T the total number of time points, and *R* the number of replicates at each time point. In our simulation studies, we fixed *p* = 2000, and considered *T* = 5,15, *C* = 5,15, and *R* = 5,15. For each setting, we repeated the simulations for 40 times.

### 3.1.2 TGCN improves the time point specific gene-gene covariance matrices estimation

The statistical nature of a gene coexpression network is a covariance matrix. Thus we first compared TGCN with the naive method in terms of the Frobenius loss in covariance matrix estimation (Fig. 1). We found that TGCN outperformed the naive methods in all simulation settings. In particular, the comparative advantage of TCGN is bigger when the number of replicates per time point decreases (e.g., R = 5) as TCGN enables the time specific covariance matrix estimates use the information from the other time points through the low-rank component. The thresholded sparse component of TCGN also effectively de-noises the estimates by trimming the spurious correlations, which could explain the more significant contrast between TCGN and the Naive method when the true covariance matrices have more complex structure (e.g., *C* = 15).

**Figure 1:**
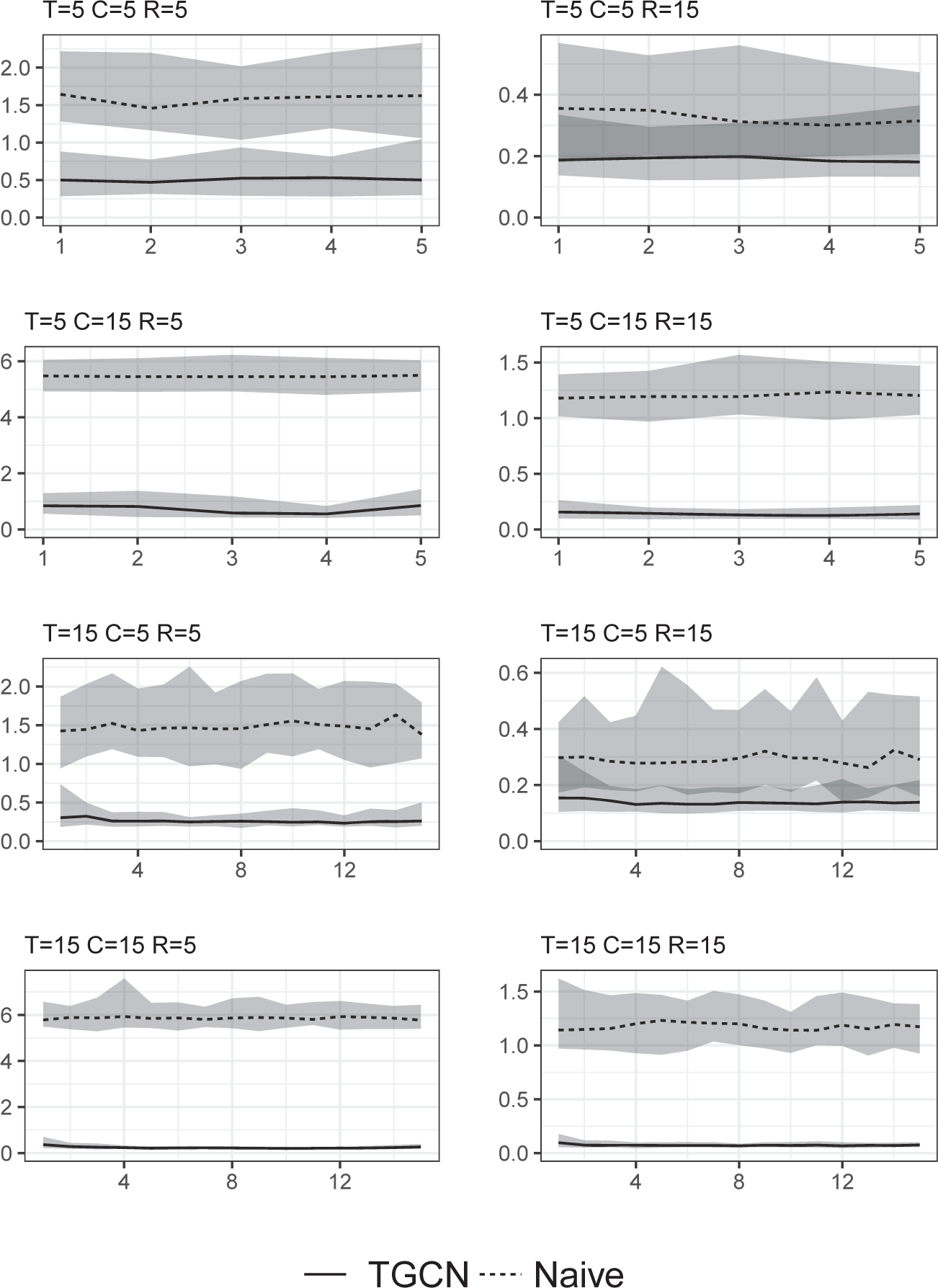
Mean Frobenius loss in covariance matrix estimation at each time point with 90% prediction band in simulations. The y axis represents the values after being normalized by the Frobenius norm of their corresponding true covariance matrices.

### 3.1.3 TGCN achieves more accurate module discovery

One of the major goals of gene coexpression analysis is module discovery, whose performance could be measured by the Adjusted Rand Index (ARI) of the discovered modules and the true module membership. In our simulation studies, the “groups” defined in the simulation setting are time-invariant and overlapping, and their group cohesiveness can also vary across time points. Thus it is not a suitable measure of the time-varying gene module architect. Instead, we used the WGCNA modules identified from the simulated true time-specific correlation matrices as the “ground truth”. Fig. 2 compared the modules discovered from the TGCN and the Naive estimates of the time-specific correlation matrices in terms of their ARI’s against the “ground truth”, and we found that TGCN led to more accurate modules estimated in most settings. Similar to our observation in the comparison in covariance matrix estimation, TGCN became more preferable when the structure of the true covariance matrices were more complex. But the number of replicates did not appear to have a clear impact on the difference between the two method, except in the case where *T* = 15 and *C* = 15.

**Figure 2:**
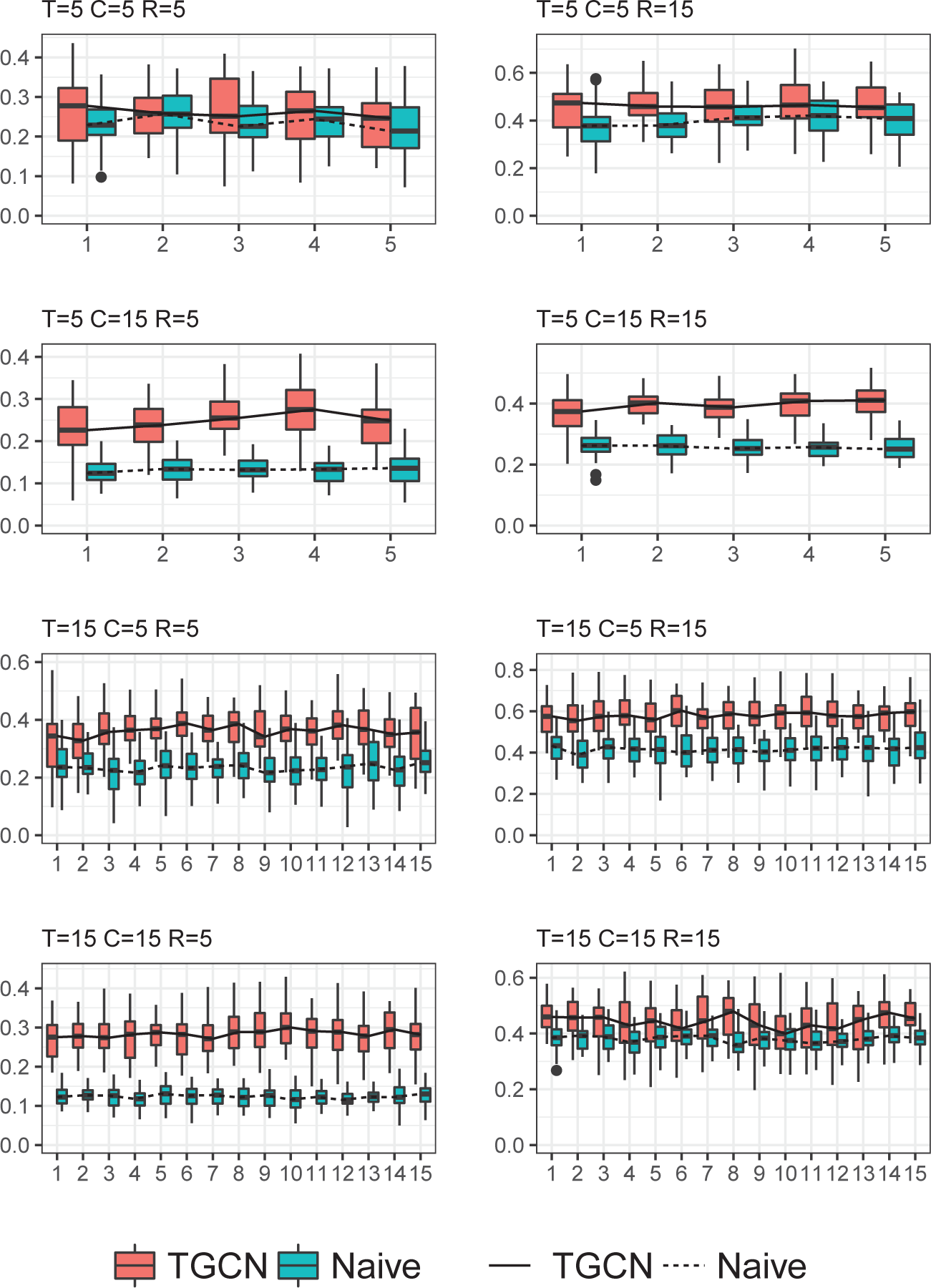
Adjusted Rand Index between the discovered modules and the “true” simulated modules at each time point.

## 3.2 Analysis of BrainSpan RNA-Seq data

### 3.2.1 Data description and preprocessing

We analyzed the RNA-Seq data from the BrainSpan Atlas of the Developing Human Brain dataset (Hawrylycz *et al*., 2012). It consists of 524 brain samples in total, grouped into 12 developmental stages ranging from eight post-conceptional weeks (pcw) to 40 years of age, including six prenatal time points and six time points after birth (see also Supplementary Table 1 for details). The numbers of samples in each stage range from 22 to 93. The expression values of this dataset were RNA-sequencing reads in the units of Per Kilobase of transcript per Million (RPKM) for 52376 genes in total. For the procedure of preprocessing the Brainspan RNA-Seq data, a log transformation (log_2_(RPKM + 1)) was applied on the expression values. We filtered out the genes with low variation in expression and consistently low expressions in the proceeding of development. Specifically, the genes with third quartile value less than log_2_(5) and the interquartile range less than log_2_(1.5) are filtered out. After the first step filtration, 10199 genes were left. For illustration purpose and to simplify the biological interpretation, we only used the 3114 genes that are involved in KEGG pathway database (Kanehisa and Goto, 2000), and consequently focused on KEGG enrichment analysis for biological interpretation. Since our goal is covariance matrix estimation, the genes expressions are centered at each time point.

### 3.2.2 TGCN extracted interpretable latent factors

In the first step of TGCN, time-invariant latent factors were learned from the normalized gene expression data with the hope that it may represent the gene group structure. We investigated the biological meanings of the 31 extracted components from the BrainSpan data in this section.

Recall that each time-invariant factor k is associated with a time-varying weight curve (*d*_*tl*_,*k*,…, *d*_*tT*_,*k)* representing the importance of this factor to the covariance matrices. Thus the factors with similar weight curves should share similar functions. We then clustered all the normalized weight curves by hierarchical clustering, and found that they fell into three clusters (Fig. 3A), each of which featured a peak around early infancy, a monotone decreasing trend, and a peak at early prenatal state, respectively. The latent factors were then grouped based on the cluster assignment.

**Figure 3:**
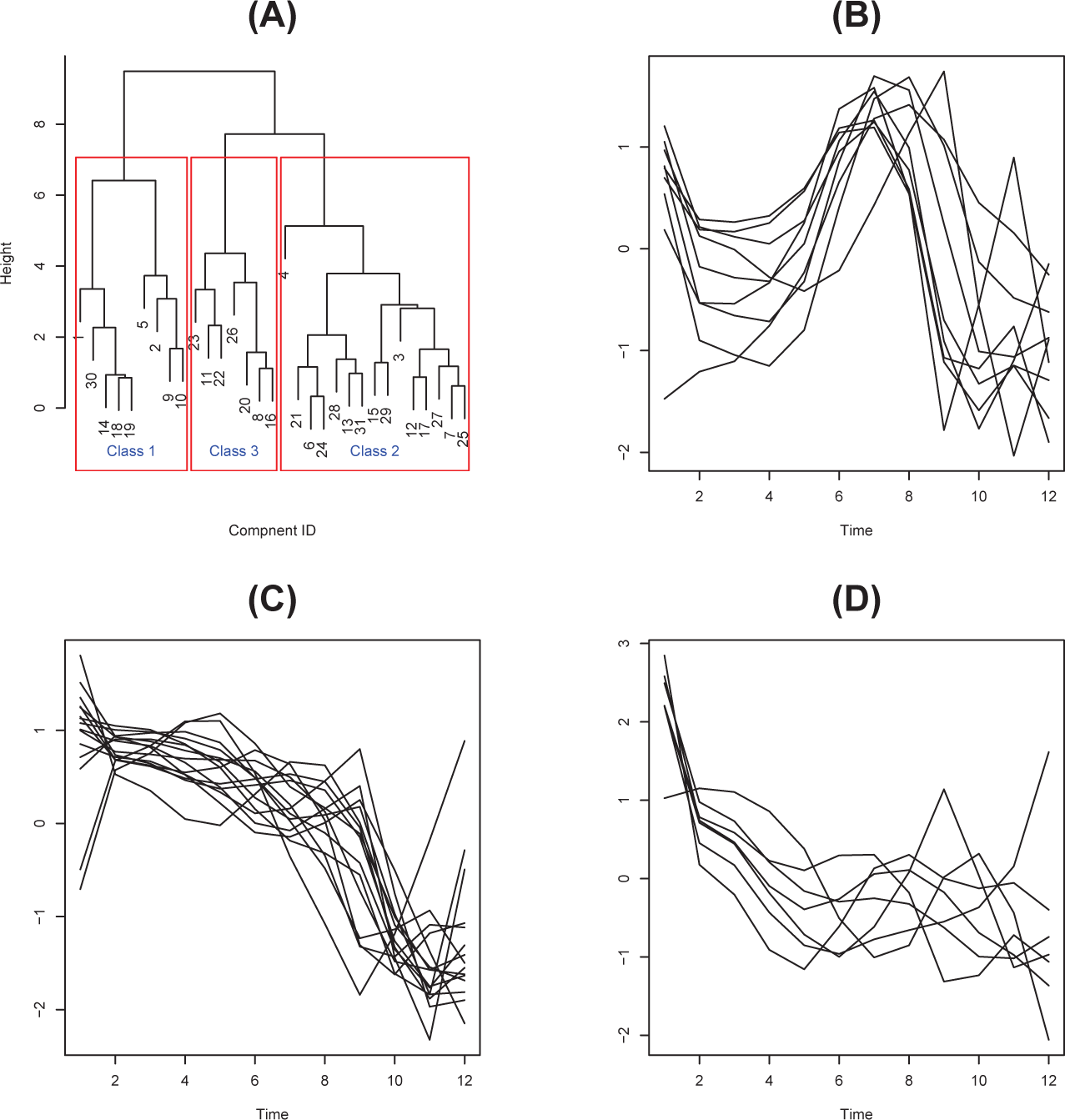
Hierarchical clustering of the normalized weight curves of the 31 TCGN factors. (A) Dendrogram of the hierarchical clustering. (B)-(D) The weight curves in the three clusters, each featured (B) a peak around early infancy, (C) a monotone decreasing trend, and (D) a peak at early prenatal state, respectively.

For each of these factor groups, we investigated the KEGG enrichment of the top 25% most relevant genes (See Supplementary Methods for details), and found that the enriched pathways were generally consistent with the overall pattern of the weight curves in each cluster. Cluster 1 featured a peak around early infancy (Fig. 3B), and the pathways enriched were generally relevant to brain development of infants. For example, Synaptic vesicle cycle pathway regulates the dendritic and synaptic density, which has been shown to reach peak in infancy and early childhood and decline from 2 to 16 yrs (Dehm *et al.*, 1972). PPR pathway is associated with white-matter development and modulates brain development in preterm infants (Krishnan *et al*., 2017; Gilmore *et al*., 2018). Cluster 2 had monotone decreasing weight curves (Fig. 3C), and was enriched with the pathways becoming inactive during aging, such as Neuroactive ligand-receptor interaction (Donerta§ *et al*., 2017). Cluster 3 contained a peak at the early prenatal state (Fig. 3D), and partially resembled the pathways relevant to the embryonic brain development such as Cell adhesion molecules (CAMs), Gap junction, and Mucin type O-glycan biosynthesis (Tran and Ten Hagen, 2013).

### 3.2.3 Module discovery and annotation

Module identification and comparison are the most widely used downstream analysis of GCN, as they reveal the potential co-regulation relationship among genes. In this section, we explored the biological interpretation of the modules discovered from the time-specific coexpression networks constructed by TGCN.

We first investigated the module conservation across time points. A new adjacency matrix was built whose edge weights were the proportion of total time points that this pair of genes were in the same module. We called a gene-gene interaction to be time-invariant if they were always in the same module. There were 1156 genes involved in such time-invariant connections, which led to a reduced adjacency matrix. Clustering based on its associated TOM distance matrix yielded 14 modules. KEGG enrichment of these modules showed that regulation of actin cytoskeleton, Protein processing in endoplasmic reticulum (ER), Ras signaling pathway, MAPK signaling pathway, and Rap1 signaling pathway were enriched (FDR = 0.1 for each module). The regulation of the actin cytoskeleton pathway is critical for the development of neural system, especially for neuronal migration (Solecki *et al*., 2009). Endoplasmic reticulum is related to various acute disorders and degenerative diseases of the brain (Paschen, 2003). The Ras and MAPK signaling pathways regulate many cell functions such as cell proliferation, survival and apoptosis. Rap1 pathway is important for Neuronal Progenitor Cell Differentiation (Rueda *et al*., 2002). Overall, these pathways encode fundamental cell functions that are expected to have strong effects at all time points, which could explain why these genes were always connected.

Differential network analysis is a popular downstream analysis after gene network construction. Thus we designed a conservative differential analysis of the pathways enriched in modules discovered at different time points (Supplementary Notes). In our study, we compared the TGCN outputs of the second (10-12 weeks prenatal) and the 11th (adolescence) time points based on this method as a showcase. It was striking to see that Huntington’s disease was enriched in two modules for the adolescence stage. Symptoms of Huntington’s disease usually begin between 30 and 50 years of age (Walker, 2007). Our results suggested that its molecular signature could be found in transcriptomic data at an even earlier age. On the other hand, the pathway enriched at the 10-12 weeks prenatal stage was Adherens junction, which was “important for maintaining tissue architecture and cell polarity and can limit cell movement and proliferation” (Kanehisa and Goto, 2000).

### 3.2.4 TGCN yields more robust gene modules in real data analysis

We also evaluated the robustness of the TGCN module output via sub-sampling. Specifically, we half-sampled the original data at each time points and ran TGCN, which repeated for 20 times. Fig. 4A compared the TGCN and the Naive method in terms of the ARI between the gene module output of the half-sampled data with the original data output. We found that TGCN yielded more consistent ARI across time points, and they were higher than the results of naive method at the majority of the time points. We remark that the time difference between the last four times points are much larger than those between the earlier time points and the time span for each of these groups are also much wider (See Supplementary Table 1), which could potentially explain the vanishing advantage of TCGN at these time points as the information of the other time points became less useful. Nevertheless, TGCN provided overall more robust modules. We also ran a real data driven simulation by adding white noise to the data. The standard deviation of the added white noise for each gene at each time point is 0.1 times the standard deviation of its expression. In this analysis, we again found that the ARI of TGCN output is higher than that of the naive estimate (Fig. 4B).

**Figure 4:**
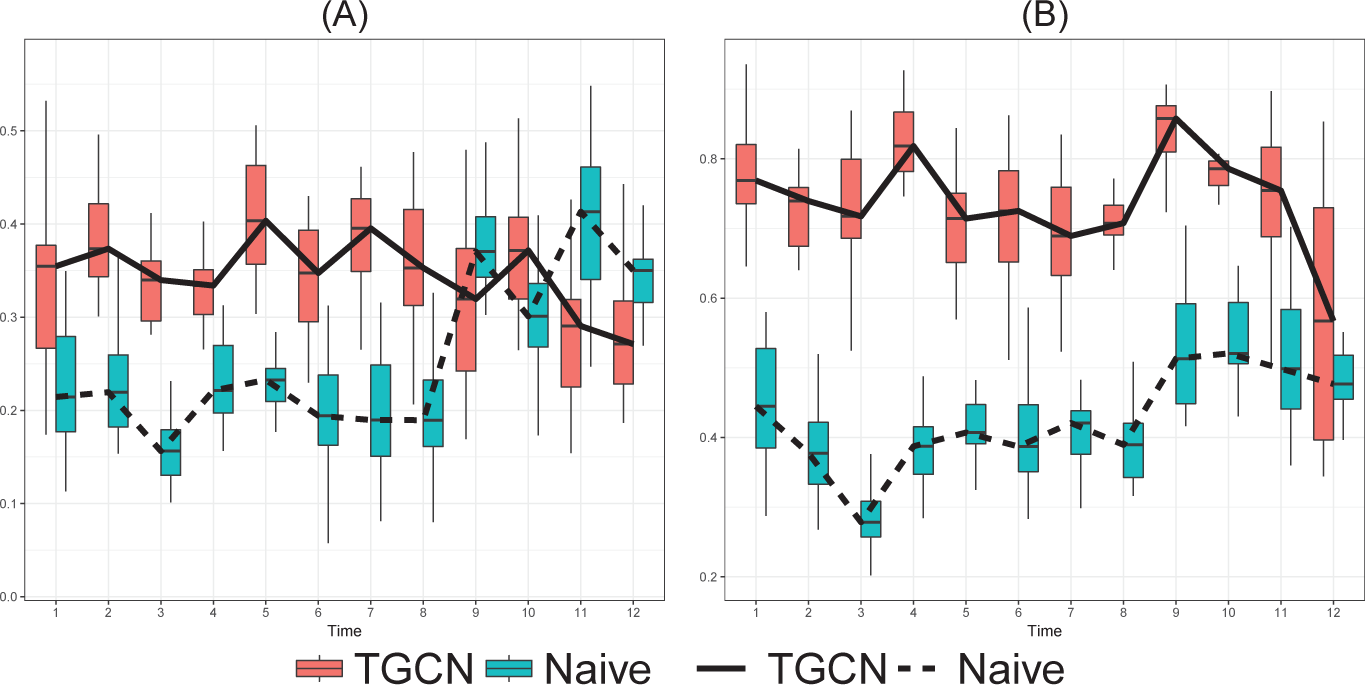
Adjusted Rand Index for (A) half-sampling the real data and (B) the real data driven simulation with 10% added noise.

## 4 Discussion

Both of gene coexpression network analysis and temporal transcriptomic studies have been widely used. There has not been any appropriate and computationally feasible time-specific GCN inference from time-course data. Most of existing studies either model the GCN at each time point completely separately, or pool the data across time points to build one single network. In this paper, we proposed Temporal Gene Coexpression Network (TGCN) that jointly model the temporal transcriptomic data when the samples at different time points are from distinct subjects. The outputs of TGCN are time-specific gene-gene correlation matrices, which allows the users to perform various downstream analysis flexibly using other computational tools such as WGCNA. Using both simulation and real data examples, we have shown that TCGN achieves more accurate correlation matrix estimation and more robust module identification.

The statistical nature of TGCN is a “low-rank plus sparse” estimator of the covariance matrices, and it could be viewed as an extension of the Principal Orthogonal Complement Thresholding (POET, Fan *et al.* 2013) method. While POET focused on one single covariance matrix, TCGN jointly estimates multiple covariance matrices simultaneously under the structural assumption that their low-rank components share the same eigen vectors.

In this paper, we focused on the correlation based gene network, as it is the most plausible model choice for the small sizes. One issue with TGCN, along with other correlation based network models, is the interpretability of the network links, as there are many biological and technical factors that may contribute to the empirical gene-gene correlations. Besides improving the data pre-processing to reduce the impact of the undesirable factors, we will also study more comprehensive models for temporal gene network analysis in the future. For example, there have been intensive statistical literature in the joint estimation of multiple gaussian graphical models (GGM) with similar structures (Peterson *et al*., 2015; Qiu *et al*., 2016; Danaher *et al*., 2014). These methods typically require relatively large sample sizes. It would be interesting to explore the possibility of extending our idea to GGM estimation. Another possible direction of future research is improving the biological intepretability of gene networks via data integration (Chen *et al*., 2017). We will explore the incorporation of the epigenetic data, metabolic pathway, gene oncology and protein-protein interactions in our modeling framework.

## Acknowledgements

This work has been supported by NSF DBI-1564621 to Drs. Chi Zhang and Qi Zhang.

## Supplementary Materials for “Temporal Gene Coexpression Network Analysis Using A Low-rank plus Sparse Framework”

## Supplementary Notes

### AIC and MDL for selecting *K*

When the number of principal components taking the value of *k* = 1, …*m*, AIC and MDL have the form of

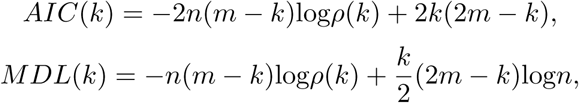

respectively, where *m* = *n* — *n*_*t*_, the function *p*(*k*) is

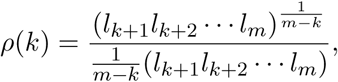

and *I*_*k*_ represents the corresponding eigenvalues 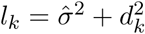 *d_k_* is *k^th^*singular value, and the value of 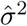 is estimated as the mean of diagonal elements of the covariance matrix after subtracting the low-rank component *UDU*^T^.

### Simulation model

We simulated gene expression data whose covariance matrices *Σ*_*t*_ were generated based on Equation (1).

Let *p* be the number of genes and *C* the number of latent factors. We first simulated the *p* × *C* time-invariant latent factor matrix *U* as the following. As we have discussed, our model was motivated by the observation that the genes usually belong to certain functional groups with time-varying effects, and these groups may overlap. Thus we first simulated a *p* × *C* binary group membership matrix *S*, where *S(j,g)* = 1{gene *j* belong to group *g*}. For each group, we picked a random integer between [1/C, 2/C] as the group size, and its members are randomly selected. Note that its rows could contain more than one non-zero elements as the groups could overlap. Then we used the left singular vectors of *S* + 0.01*A* as the latent factors *U*. Here *A* is a *p* × *C* random matrices whose entries are i.i.d. from *Unif* (—1,1).

Next, we simulated the time-varying weights of these latent factors. For *t* = 1,…, *T*, let *d_g_,_t_* be the *g*th diagonal element of *D*_*t*_. We defined (log(*d*_*g,1*_),…, log(*d*_*g,T*_)) as a random linear combination of B-spline basis defined on (0,*T* + 1), whose coefficients were i.i.d. samples from *Unif* (—1,1). Then *L*_*t*_ = *U D_t_U^T^* is the low-rank component of the simulated covariance matrix.

The most naive way of simulating the sparse component *R*_*t*_ is simply generating a sparse random symmetric matrix. But such matrix does not contain any information about the structure. To generate an informative sparse component, we defined the elements of the upper triangle of the symmetric matrix *R*_*t*_ as the following.

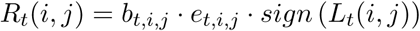

where 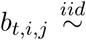 *Unif* (0.1,0.3), 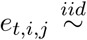 *Bernoulli*(*p*_*t,i,j*_) where *p*_*t,i,j*_ = 0.005 · 1{|*L*_*t*_(*i,j*)| > *q*_*t*_} + 0.0005 · 1{|*L*_*t*_(*i, j*)| < *q*_*t*_}, and the diagonal elements of *R*_*t*_ were set to 1. Here *b*_*i,j*_ modeled the magnitude of time-point specific sparse component, and the definition of *p*_*t,i,j*_ ensured that *R*_*t*_ and *L*_*t*_ contained non-contradicting information about the underlying structure. In our simulation study, we set *q*_*t*_ as the 0.9 quantile of the absolute values of the off-diagonal elements of *L*_*t*_.

Finally, we defined the time-specific covariance matrix *Σ*_*t*_ = *U D_t_U^T^* + *R*_*t*_, and simulated *X_t_,_r_* from N(0, Σ_*t*_ for *r* = 1,…, *R*. In our simulation studies, we fixed *p* = 2000, and considered *T* = 5,15, *C* = 5,15, and *R* = 5,15. For each setting, we repeated the simulations for 40 times.

### Identifying the most relevant genes for latent factors

According to Equation (1), for each pair of genes (*i, j*),

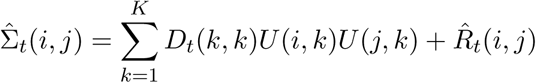

The contribution of one single factor k to the correlation pattern of gene i to the other genes could be measured by |*U*(*i, k*)|, because factor k becomes irrelevant to the covariances involving gene i at all time points if |*U*(*i,k*)| is close to 0. Following a similar spirit, the joint relevance of a group of factors *k*_1_,…,*k*_*A*_ to the covariances involving a particular gene *i* could be measured by 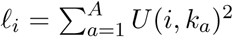. It is essentially the *L*_2_ row norm of the corresponding submatrix of *U*. For a group of factors, we said a gene *i* was more relevant to these factors if l_*i*_ is large. For our KEGG enrichment analysis, we considered the top 25% genes with the largest *l*_*i*_ for each cluster of latent factors.

### Conservative differential enrichment analysis

In a pairwise comparison of time points *t***1** and *t***2**, a pathway is said to be specific to *t***1** if it satisfies the following conditions. (1) It is enriched for some modules at *t*_1_ but not at *t*_2_ nor for the above time-invariant modules; and (2) at most 25% of the genes that are involved in this pathway are also in any of the pathways enriched at t**2** or in the time-invariant modules. The second criterion is to avoid the case where two pathways shared a large proportion of genes, but were enriched at different time points due to their minor differences in gene composition and the statistical cutoff in enrichment analysis.

## Supplementary Figures

**Figure 1:**
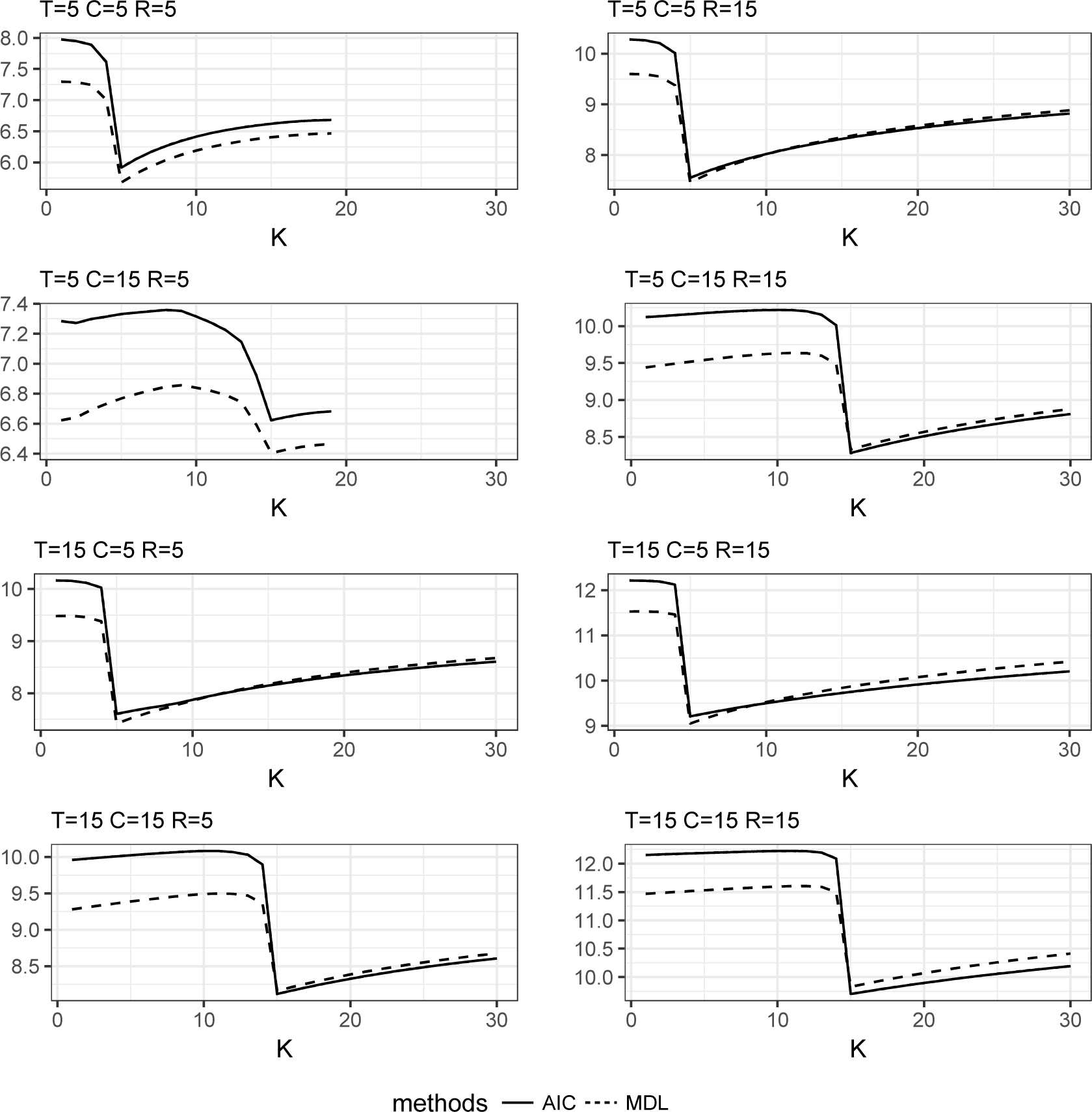
Selecting the number of latent factors using AIC and MDL for the simulation studies

**Figure 2:**
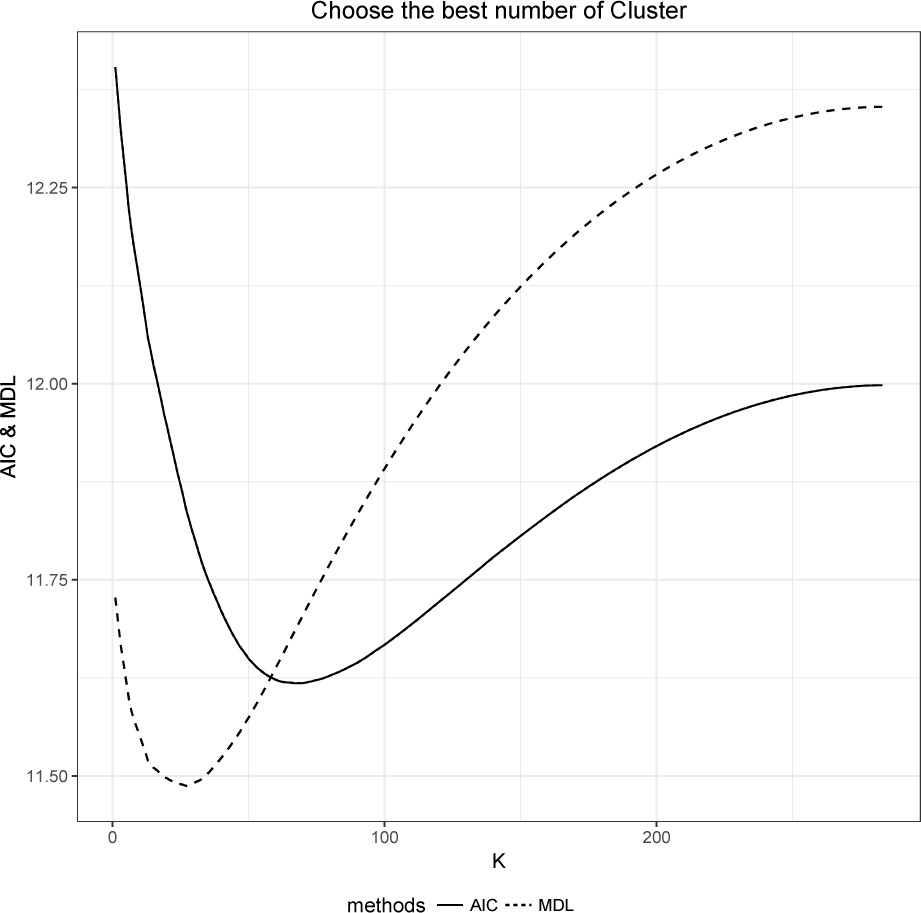
Selecting the number of latent factors using AIC and MDL for the real data analysis

## Supplementary Tables

**Table 1:** Description of developmental stages for the Brainspan data (Hawrylycz *et al.*, 2012)

| Stage | Age | Developmental stage | Replicates number |
| --- | --- | --- | --- |
| 1 | 8-9 pcw | Early prenatal | 30 |
| 2 | 10-12 pcw | Early prenatal | 45 |
| 3 | 13-15 pcw | Early mid-prenatal | 44 |
| 4 | 16-18 pcw | Early mid-prenatal | 53 |
| 5 | 19-24 pcw | Late mid-prenatal | 43 |
| 6 | 25-38 pcw | Late prenatal | 22 |
| 7 | Birth-5 months | Early infancy | 33 |
| 8 | 6-18 months | Late infancy | 26 |
| 9 | 19 months-5 yrs | Early childhood | 44 |
| 10 | 6-11 yrs | Late childhood | 41 |
| 11 | 12-19 yrs | Adolescence | 50 |
| 12 | 20-40 yrs | Adulthood | 93 |

## References

Akaike, H. (1973). Maximum likelihood identification of gaussian autoregressive moving average models. Biometrika, 60(2). 255–265.

Akaike, H. (1998). Information theory and an extension of the maximum likelihood principle. In Selected Papers of Hirotugu Akaike, pages 199–213. Springer.

Azevedo, H., Khaled, N. A., Santos, P., Bertonha, F. B., and Moreira-Filho, C. A. (2017). Temporal analysis of hippocampal ca3 gene co-expression networks in a rat model of febrile seizures. Disease Models & Mechanisms.

Ballouz, S., Verleyen, W., and Gillis, J. (2015). Guidance for rna-seq co-expression network construction and analysis: safety in numbers. Bioinformatics, 31(13), 2123–2130.

Cai, T. and Liu, W. (2011). Adaptive thresholding for sparse covariance matrix estimation. Journal of the American Statistical Association, 106(494), 672–684.

Chen, X., Gu, J., Wang, X., Jung, J.-G., Wang, T.-L., Hilakivi-Clarke, L., Clarke, R., and Xuan, J. (2017). Crnet: An efficient sampling approach to infer functional regulatory networks by integrating large-scale chip-seq and time-course rna-seq data. Bioinformatics, page btx827.

Danaher, P., Wang, P., and Witten, D. M. (2014). The joint graphical lasso for inverse covariance estimation across multiple classes. Journal of the Royal Statistical Society: Series B (Statistical Methodology), 76(2), 373–397.

Daub, C. O., Steuer, R., Selbig, J., and Kloska, S. (2004). Estimating mutual information using b-spline functions-an improved similarity measure for analysing gene expression data. BMC bioinformatics, 5(1), 118.

Dehm, P., Jimenez, S. A., Olsen, B. R., and Prockop, D. J. (1972). A transport form of collagen from embryonic tendon: electron microscopic demonstration of an nh2-terminal extension and evidence suggesting the presence of cystine in the molecule. Proceedings of the National Academy of Sciences, 69(1), 60–64.

Donertas, H. M., Izgi, H., Kamacioglu, A., He, Z., Khaitovich, P., and Somel, M. (2017). Gene expression reversal toward pre-adult levels in the aging human brain and age-related loss of cellular identity. Scientific reports, 7(1), 5894.

Fan, J., Fan, Y., and Lv, J. (2008). High dimensional covariance matrix estimation using a factor model. Journal of Econometrics, 147(1), 186–197.

Fan, J., Liao, Y., and Mincheva, M. (2013). Large covariance estimation by thresholding principal orthogonal complements. Journal of the Royal Statistical Society: Series B (Statistical Methodology), 75(4), 603–680.

Ficklin, S. P., Luo, F., and Feltus, F. A. (2010). The association of multiple interacting genes with specific phenotypes in rice using gene coexpression networks. Plant physiology, 154(1), 13–24.

Friedman, N., Linial, M., Nachman, I., and Pe’er, D. (2000). Using bayesian networks to analyze expression data. Journal of computational biology, 7(3–4), 601–620.

Gao, C., McDowell, I. C., Zhao, S., Brown, C. D., and Engelhardt, B. E. (2016). Context specific and differential gene co-expression networks via bayesian biclustering. PLoS computational biology, 12(7), e1004791.

Gilmore, J. H., Knickmeyer, R. C., and Gao, W. (2018). Imaging structural and functional brain development in early childhood. Nature Reviews Neuroscience, 19(3), 123.

Hawrylycz, M. J., Lein, S., Guillozet-Bongaarts, A. L., Shen, E. H., Ng, L., Miller, J. A., Van De Lagemaat, L. N., Smith, K. A., Ebbert, A., Riley, Z. L., et al. (2012). An anatomically comprehensive atlas of the adult human brain transcriptome. Nature, 489(7416), 391.

Kadarmideen, H. N. and Watson-Haigh, N. S. (2012). Building gene co-expression networks using transcriptomics data for systems biology investigations: Comparison of methods using microarray data. Bioinformation, 8(18), 855.

Kanehisa, M. and Goto, S. (2000). Kegg: kyoto encyclopedia of genes and genomes. Nucleic acids research, 28(1), 27–30.

Krishnan, M. L., Wang, Z., Aljabar, P., Ball, G., Mirza, G., Saxena, A., Counsell, S. J., Hajnal, J. V., Montana, G., and Edwards, A. D. (2017). Machine learning shows association between genetic variability in pparg and cerebral connectivity in preterm infants. Proceedings of the National Academy of Sciences, page 201704907.

Langfelder, P. and Horvath, S. (2008). Wgcna: an r package for weighted correlation network analysis. BMC bioinformatics, 9(1), 559.

Lebre, S., Becq, J., Devaux, F., Stumpf, M. P., and Lelandais, G. (2010). Statistical inference of the time-varying structure of gene-regulation networks. BMC systems biology, 4(1), 130.

Luo, X. (2011). High dimensional low rank and sparse covariance matrix estimation via convex minimization. Arxiv preprint.

Miao, Z., Han, Z., Zhang, T., Chen, S., and Ma, C. (2017). A systems approach to a spatio-temporal understanding of the drought stress response in maize. Scientific reports, 7(1), 6590.

Ni, Y., Stingo, F. C., and Baladandayuthapani, V. (2015). Bayesian nonlinear model selection for gene regulatory networks. Biometrics, 71(3), 585–595.

Oates, C. J. and Mukherjee, S. (2012). Network inference and biological dynamics. The annals of applied statistics, 6(3), 1209.

Obayashi, T., Hayashi, S., Shibaoka, M., Saeki, M., Ohta, H., and Kinoshita, K. (2007). Coxpresdb: a database of coexpressed gene networks in mammals. Nucleic acids research, 36(suppl_1), D77–D82.

Paschen, W. (2003). Endoplasmic reticulum: a primary target in various acute disorders and degenerative diseases of the brain. Cell calcium, 34(4–5), 365–383.

Peterson, C., Stingo, F. C., and Vannucci, M. (2015). Bayesian inference of multiple gaussian graphical models. Journal of the American Statistical Association, 110(509), 159–174.

Qiu, H., Han, F., Liu, H., and Caffo, B. (2016). Joint estimation of multiple graphical models from high dimensional time series. Journal of the Royal Statistical Society: Series B (Statistical Methodology), 78(2), 487–504.

Rissanen, J. (1978). Modeling by shortest data description. Automatica, 14(5), 465–471.

Rissanen, J. (1983). A universal prior for integers and estimation by minimum description length. The Annals of statistics, pages 416–431.

Rueda, D., Navarro, B., Martinez-Serrano, A., Guzman, M., and Galve-Roperh, I. (2002). The endocannabinoid anandamide inhibits neuronal progenitor cell differentiation through attenuation of the rap1/b-raf/erk pathway. Journal of Biological Chemistry, 277(48), 46645–46650.

Schafer, J. and Strimmer, K. (2004). An empirical bayes approach to inferring large-scale gene association networks. Bioinformatics, 21(6), 754–764.

Solecki, D. J., Trivedi, N., Govek, E.-E., Kerekes, R. A., Gleason, S. S., and Hatten, M. E. (2009). Myosin ii motors and f-actin dynamics drive the coordinated movement of the centrosome and soma during cns glial-guided neuronal migration. Neuron, 63(1), 63–80.

Song, L., Langfelder, P., and Horvath, S. (2012). Comparison of co-expression measures: mutual information, correlation, and model based indices. BMC bioinformatics, 13(1), 328.

Tan, M., Cheng, D., Yang, Y., Zhang, G., Qin, M., Chen, J., Chen, Y., and Jiang, M. (2017). Co-expression network analysis of the transcriptomes of rice roots exposed to various cadmium stresses reveals universal cadmium-responsive genes. BMC plant biology, 17(1), 194.

Tran, D. T. and Ten Hagen, K. G. (2013). Mucin-type o-glycosylation during development. Journal of Biological Chemistry, 288(10), 6921–6929.

Valle, S., Li, W., and Qin, S. J. (1999). Selection of the number of principal components: the variance of the reconstruction error criterion with a comparison to other methods. Industrial & Engineering Chemistry Research, 38(11), 4389–4401.

van Dam, S., Craig, T., and de Magalhaes, J. P. (2014). Genefriends: a human rna-seq-based gene and transcript co-expression database. Nucleic acids research, 43(D1), D1124–D1132.

Villa-Vialaneix, N., Liaubet, L., Laurent, T., Cherel, P., Gamot, A., and SanCristobal, M. (2013). The structure of a gene co-expression network reveals biological functions underlying eqtls. PloS one, 8(4), e60045.

Walker, F. O. (2007). Huntington’s disease. The Lancet, 369(9557), 218–228.

Xu, H., Gao, Y., and Wang, J. (2012). Transcriptomic analysis of rice (oryza sativa) developing embryos using the rna-seq technique. PLoS One, 7(2), e30646.

Yang, Y., Han, L., Yuan, Y., Li, J., Hei, N., and Liang, H. (2014). Gene co-expression network analysis reveals common system-level properties of prognostic genes across cancer types. Nature communications, 5, 3231.

Yim, W. C., Yu, Y., Song, K., Jang, C. S., and Lee, B.-M. (2013). Planex: the plant co-expression database. BMC plant biology, 13(1), 83.

Yu, G., Wang, L.-G., Han, Y., and He, Q.-Y. (2012). clusterprofiler: an r package for comparing biological themes among gene clusters. Omics: a journal of integrative biology, 16(5), 284–287.

Zhang, B. and Horvath, S. (2005). A general framework for weighted gene co-expression network analysis. Statistical applications in genetics and molecular biology, 4(1).

Zhang, Q., Yu, Y., Zhang, J., and Liang, H. (2018). Using single-index odes to study dynamic gene regulatory network. PloS one, 13(2), e0192833.

Zhu, S. and Wang, Y. (2015). Hidden markov induced dynamic bayesian network for recovering time evolving gene regulatory networks. Scientific reports, 5, 17841.

Zou, M. and Conzen, S. D. (2004). A new dynamic bayesian network (dbn) approach for identifying gene regulatory networks from time course microarray data. Bioinformatics, 21(1), 71–79.

